# Blue/green light-responsive cyanobacteriochromes are cell shade sensors in red-light replete niches

**DOI:** 10.1101/839886

**Authors:** Gen Enomoto, Masahiko Ikeuchi

## Abstract

Photoautotrophic cyanobacteria have developed sophisticated light response systems to capture and utilize the energy and information of incident light [1]. Cyanobacteria-specific photoreceptors cyanobacteriochromes (CBCRs) are distantly related to more widespread phytochromes. CBCRs show the most diverse spectral properties among the naturally occurring photoreceptors, typified by a unique and prevailing blue/green light-absorbing variant [2–6]. However, where the CBCR-mediated ‘colorful’ signaling systems function in nature has been elusive. We previously reported that the three CBCRs SesA/B/C synthesize/degrade a bacterial second messenger cyclic diguanylate (c-di-GMP) in response to blue/green light [6–8]. The cooperative action of SesA/B/C enables blue light-ON and green light-OFF regulation of the c-di-GMP-dependent cell aggregation of the thermophilic cyanobacterium *Thermosynechococcus vulcanus* [8, 9]. Here, we report that SesA/B/C can serve as a physiological sensor of cell density. Because cyanobacterial cells show lower transmittance of blue light than green light, higher cell density gives more green light-enriched irradiance to cells. The cell density-dependent suppression of cell aggregation under blue/green-mixed light and white light conditions support this idea. Such a sensing mechanism may provide information about the cell position in cyanobacterial mats in hot springs, the natural habitat of *Thermosynechococcus*. This cell position-dependent SesA/B/C-mediated regulation of cellular sessility (aggregation) might be ecophysiologically essential for the reorganization and growth of phototrophic mats. We also report that the green light-induced dispersion of cell aggregates requires red light-driven photosynthesis. Blue/green CBCRs might work as shade detectors in a different niche than red/far-red phytochromes, which may be why CBCRs have evolved in cyanobacteria.

## Results and Discussion

### The Ratio of Blue to Green Light Perceived by CBCRs SesA/B/C Can be an Indicator of How Many Other Cells Shade Them

The absorption spectrum of *Thermosynechococcus* cells shows a peak at 435 nm and a trough at ~530 nm, which roughly correspond to the blue absorption (~430 nm) and the teal-green absorption (500 ~ 530 nm) of the three cyanobacteriochromes, SesA/B/C (Figure 1A) [8]. Since white light can be differentially absorbed by cells containing chlorophyll, carotenoids and phycocyanin/allophycocyanin, the light transmitted through a given cell layer is increasingly enriched in green light, depending on cell density (Figure 1B). Thus, we hypothesized that SesA/B/C-mediated cell aggregation may be governed by cell density under natural light.

**Figure 1.**
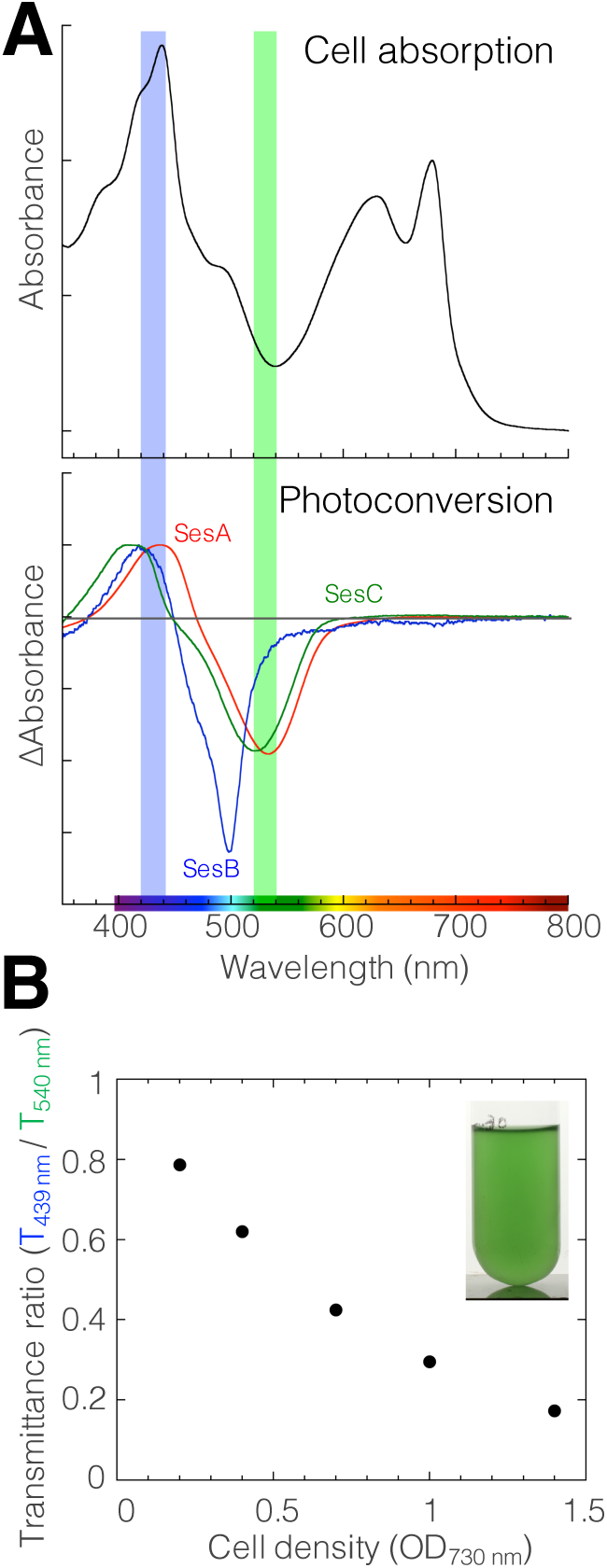
The Light Ratio of Blue to Green Light Perceived by SesA/B/C CBCRs Can be an Indicator of Self-Shading. (A) The cell absorption spectrum of *Thermosynechococcus vulcanus* (upper) and the difference spectra of the reversible photoconversion of SesA/B/C holoproteins (lower). The lower panel was created using the data reported in ref. 7. The region of 420 nm–440 nm was highlighted in a blue shade and that of 520 nm–540 nm was highlighted in a green shade. (B) The transmittance ratio of blue light (439 nm) to green light (540 nm) and the image of the liquid culture of *T. vulcanus*.

We assessed the effects of cell population density on cell aggregation under model light conditions. We used mixed light conditions of teal-green and blue (each 0~10 µmol photons m^−2^ s^−1^ as signaling light) in addition to red light for growth (30 µmol photons m^−2^ s^−1^). When ten µmol photons m^−2^ s^−1^ blue light and 2 ~ 10 µmol photons m^−2^ s^−1^ teal-green light were used as signaling light, we observed cell density-dependent suppression of cell aggregation (Figure 2A). The more green light that was provided, the less cell aggregation that was induced. Under only blue or teal-green light (10 µmol photons m^−2^ s^−1^), cell aggregation did not depend on the cell density; cells showed secure cell aggregation irrespective of the density under only blue light, whereas they showed no cell aggregation under teal-green light (Figure 2A). This result suggests that the cell density dependency can be mediated by the blue/teal-green light-sensing system SesA/B/C but not by other light-independent regulatory systems, such as quorum sensing. This assay also confirmed that the ratio of blue light/teal-green light, but not the absolute intensity of blue light, is the signal for cell aggregation, as the more teal-green light that was added to 10 µmol photons m^−2^ s^−1^ blue light, the more cell aggregation was repressed (Figure 2A).

**Figure 2.**
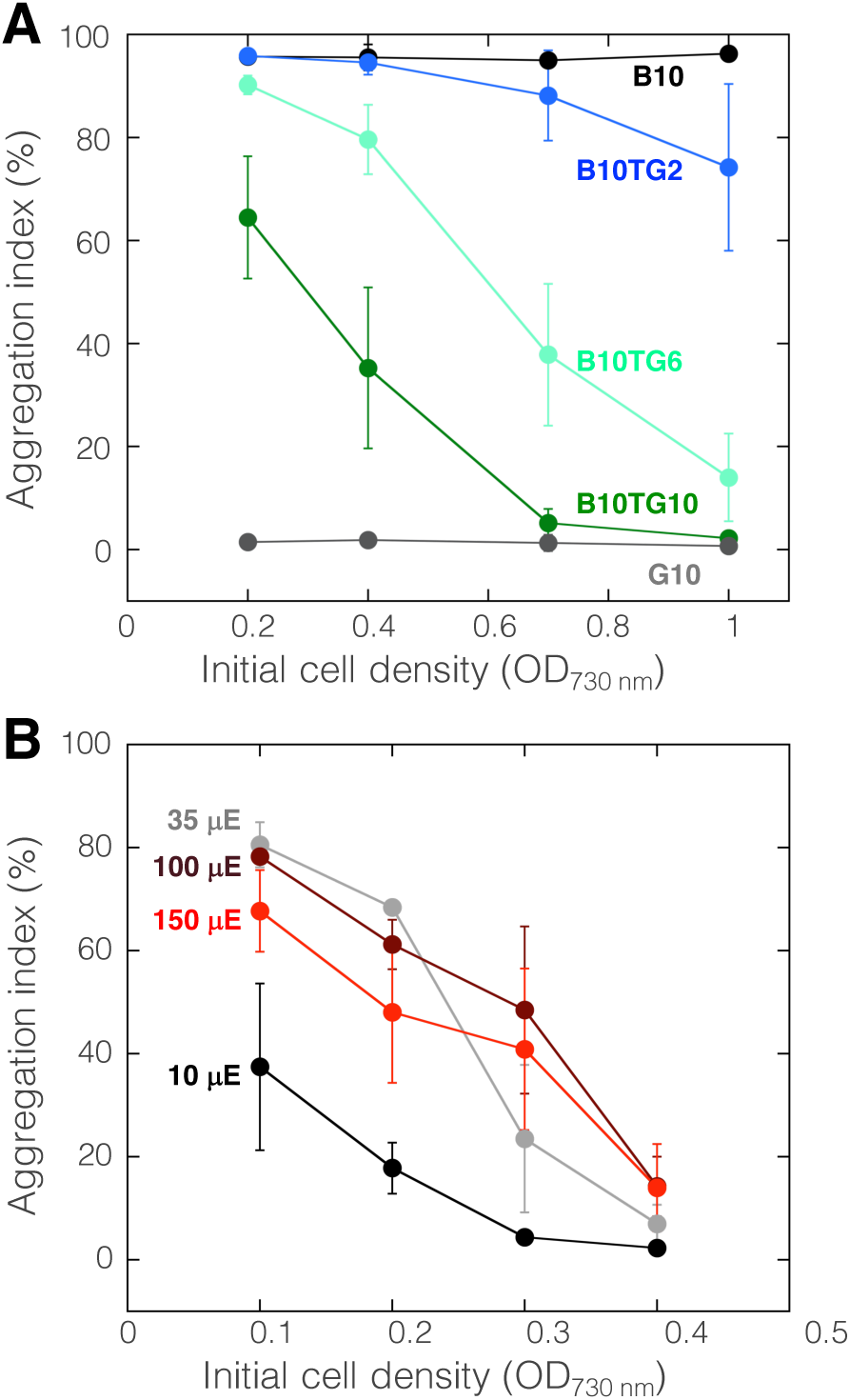
SesA/B/C-Mediated Blue/Green Light Signaling Senses Cell Density to Regulate Cell Aggregation under Both Blue Light and Green Light. (A) Effects of cell population density on cell aggregation under LED irradiation. The horizontal axis shows the initial cell density, and the vertical axis shows the aggregation index after 48 h of cultivation at low temperatures of 31°C in the presence of red light (634 nm, 30 µE m^−2^ s^−1^). The intensity of signaling light is indicated as labels; B10TG2: blue light (448 nm, 10 µE m^−2^ s^−1^) + teal-green light (507 nm, 2 µE m^−2^ s^−1^). Data are represented as the mean ± SEM. (B) Cell aggregation under the four different intensities of the white light fluorescent lamp. Data are represented as the mean ± SEM.

To extend our model light system to naturally occurring conditions, we assessed the cell density dependency of cell aggregation under white light irradiation. Under white light ranging from 10 to 150 µmol photons m^−2^ s^−1^, cell aggregation decreased as the cell density increased (Figure 2B). However, under low light of 10 µmol photons m^−2^ s^−1^, aggregation was weak even at the lowest cell density, probably because the light was insufficient for photosynthesis or activation of the SesA/B/C photoreceptors. The aggregation at the lowest cell density under high light of 150 µmol photons m^−2^ s^−1^ tended to be weaker than that under 30 or 100 µmol photons m^−2^ s^−1^. This lowered cell aggregation may be due to the high light stress because light irradiation with higher than 150 µmol photons m^−2^ s^−1^ might be dangerous for low-density cell cultures under the low-temperature condition of 31°C. These results suggest that cell density-dependent control of cell aggregation can also be mediated by the blue/teal-green light-sensing proteins SesA/B/C under white light.

### Blue/Green Light-Regulated Reversible Formation and Dispersion of Cell Aggregates are Driven by Red Light-Driven Photosynthesis

To dissect the contribution of each wavelength of light, we assessed the role of red light on blue/green light-regulated cell aggregation. When cell aggregates were transferred from blue light conditions to teal-green light conditions at 31°C with red background light, they dispersed in 24 h (Figure 3A). This result indicates that cell aggregation is reversible even under cell aggregation-enhancing low-temperature conditions. The omission of the background red light impaired the dispersion of cell aggregates (Figure 3A). These results suggested that the background red light may support dispersion via photosynthetic activity, given that photosynthetic pigments poorly absorb teal-green light. Consistently, the addition of the photosynthesis inhibitor 3-(3,4-dichlorophenyl)-1,1-dimethylurea (DCMU) impaired both the formation and dispersion of cell aggregates (Figure 3B). Thus, growth light is critical for the decision and active responses to induce a sessile lifestyle (cell aggregation) or planktonic lifestyle (dispersion). Many cyanobacteria, including *Thermosynechococcus*, possess chlorophyll *a* and phycocyanin pigments for photosynthesis but not far-red light-absorbing chlorophyll *d*/*f* or green light-absorbing phycoerythrin. This pigment composition means that blue, orange, and red light but not green or far-red light is effective for the active photosynthesis-driven responses. In other words, irradiation with only blue light can induce cell aggregation with active photosynthesis (Figure 3B). In contrast, green light should be accompanied by orange light or red light for the induction of cell dispersion. Both the formation and dispersion of cellulose-dependent cell aggregates [10] may be energy-demanding responses because cells will have to produce a massive amount of extracellular cellulose to stabilize the cell aggregates during formation and degrade the cellulose when they escape from the aggregates.

**Figure 3.**
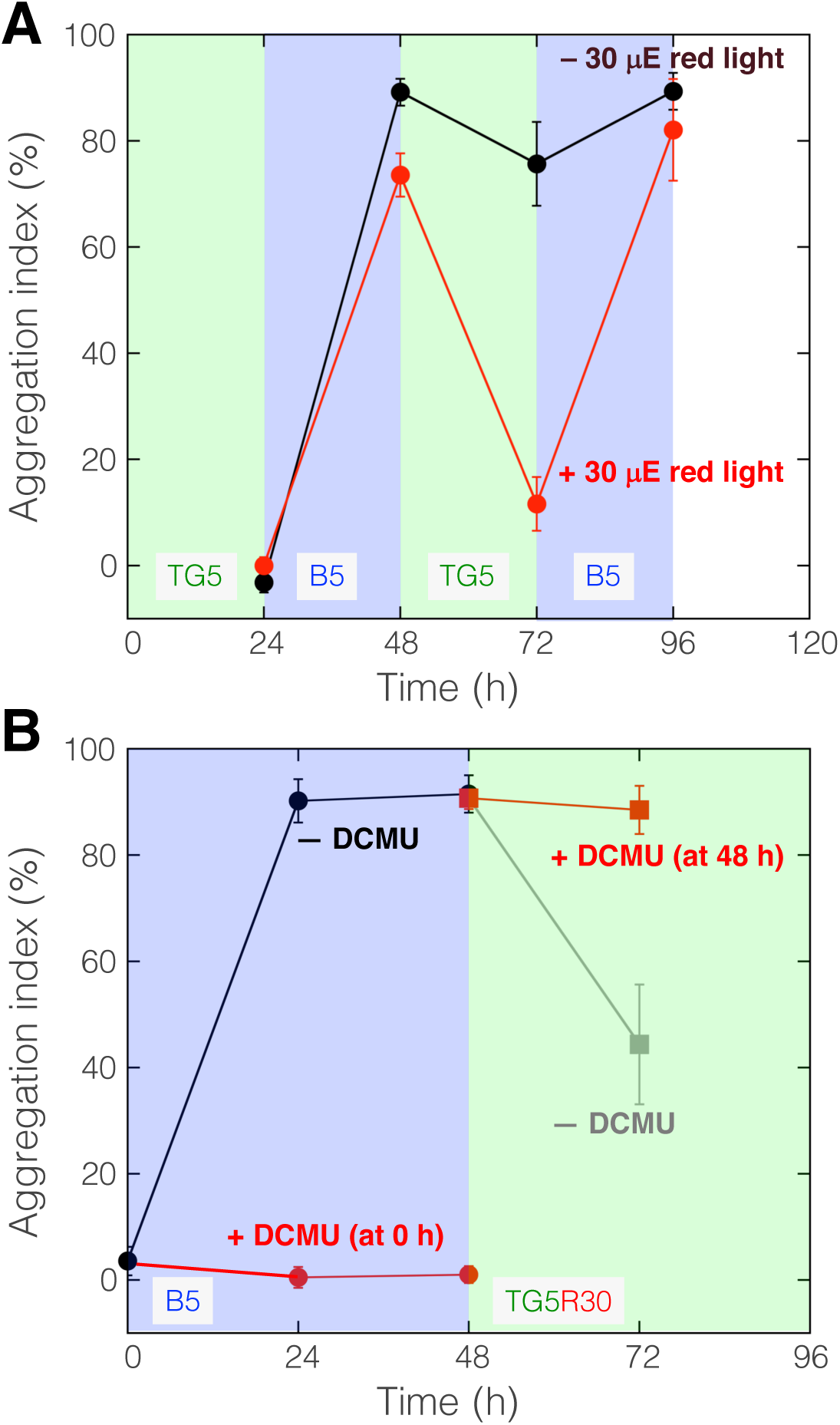
Red Light-Driven Photosynthesis is Necessary to Support the Reversible Formation and Dispersion of Cell Aggregates. (A) Reversible formation and dispersion of cell aggregation induced by switching blue light and teal-green light every 24 h in the presence or absence of the background red light irradiation. Data are represented as the mean ± SD. (B) Effects of the photosynthesis inhibitor DCMU on the formation and dispersion of cell aggregates. DCMU (final concentration 10 µM) was added at the indicated time.

### The Physiological and Ecological Relevance of Blue/Green-Type CBCR Signaling

Many bacteria often establish multicellular, matrix-embedded communities, such as microbial mats or biofilms, which are crucial for their physiology, ecology, and infections [11, 12]. Likewise, thermophilic cyanobacteria usually form a microbial mat in hot springs [13, 14]. Analysis of the transmittance of light in typical thermophilic cyanobacterial mats revealed that green and far-red light are penetrating, whereas blue and red light are mostly absorbed by photosynthetic pigments [15, 16]. This tendency fits well with the light transmittance properties of the cell suspension of *Thermosynechococcus* (Figure 1A). Thus, reversible regulation of sessility by blue/green CBCR signaling could account for the behavioral responses of *Thermosynechococcus* in the mats.

Physiologically, low temperature-enhanced, and blue light-induced cell aggregation would protect photosynthesis from photoinhibition due to self-shading effects, as suggested previously [10]. Blue light irradiation easily damages the oxygen-evolving complex of photosystem II [17]. Usually, the rapid repair cycle of photosystem II supports the replacement of the damaged reaction center polypeptides with newly synthesized ones [18]. At low temperatures, on the other hand, the protein turnover is decelerated, leading to impaired repair of the photosystem. Under these photoinhibition-inducing conditions, the formation of cell aggregates may be beneficial for the protection of the photosynthetic apparatus by self-shading. However, low temperature is not necessary to induce sessile lifestyles because blue light signals can induce cell aggregation alone, even at 45°C [19]. Because the formation of cell aggregates is enhanced at relatively low temperatures, it will be beneficial for cells to remain in the hot spring environment and to avoid being displaced by water currents to lethally low-temperature environments such as rivers and oceans.

Ecologically, light-induced regulation of sessility and motility would be necessary for the remodeling (or reorganizing) of phototrophic mats. The phototactic motility of *Thermosynechococcus* is regulated intricately by light quality and intensity [20]. c-di-GMP is often involved in the regulation of bacterial motility [21, 22]. As cell aggregates develop, the internal cells in a floc will sense a green light-rich environment and turn off c-di-GMP signaling, favoring the motile-planktonic lifestyle. Given that the *Thermosynechococcus vulcanus* strain used in this study shows positive phototaxis, the internal cells might move through the crowd of cells toward the sun. When these cells reach the upper region, they will sense the relatively blue light-rich environment and turn on c-di-GMP signaling, leading to a sessile mode of life. Then, the next generation of internal cells will sense the green light-rich environment and turn off c-di-GMP signaling. Repeating this process may lead to dynamic cell movements inside a static microbial mat (Figure 4A). This dynamism might facilitate nutrient uptake and gas exchange, which are necessary for efficient photosynthetic growth, and might also be helpful for the expansion of a microbial mat. The other potential advantage is for every cell of the community to obtain access to light for photosynthetic viability and growth. Further *in situ* and *ex situ* studies are needed to reveal the role and significance of c-di-GMP heterogeneity and cellular behaviors in a cyanobacterial community.

**Figure 4.**
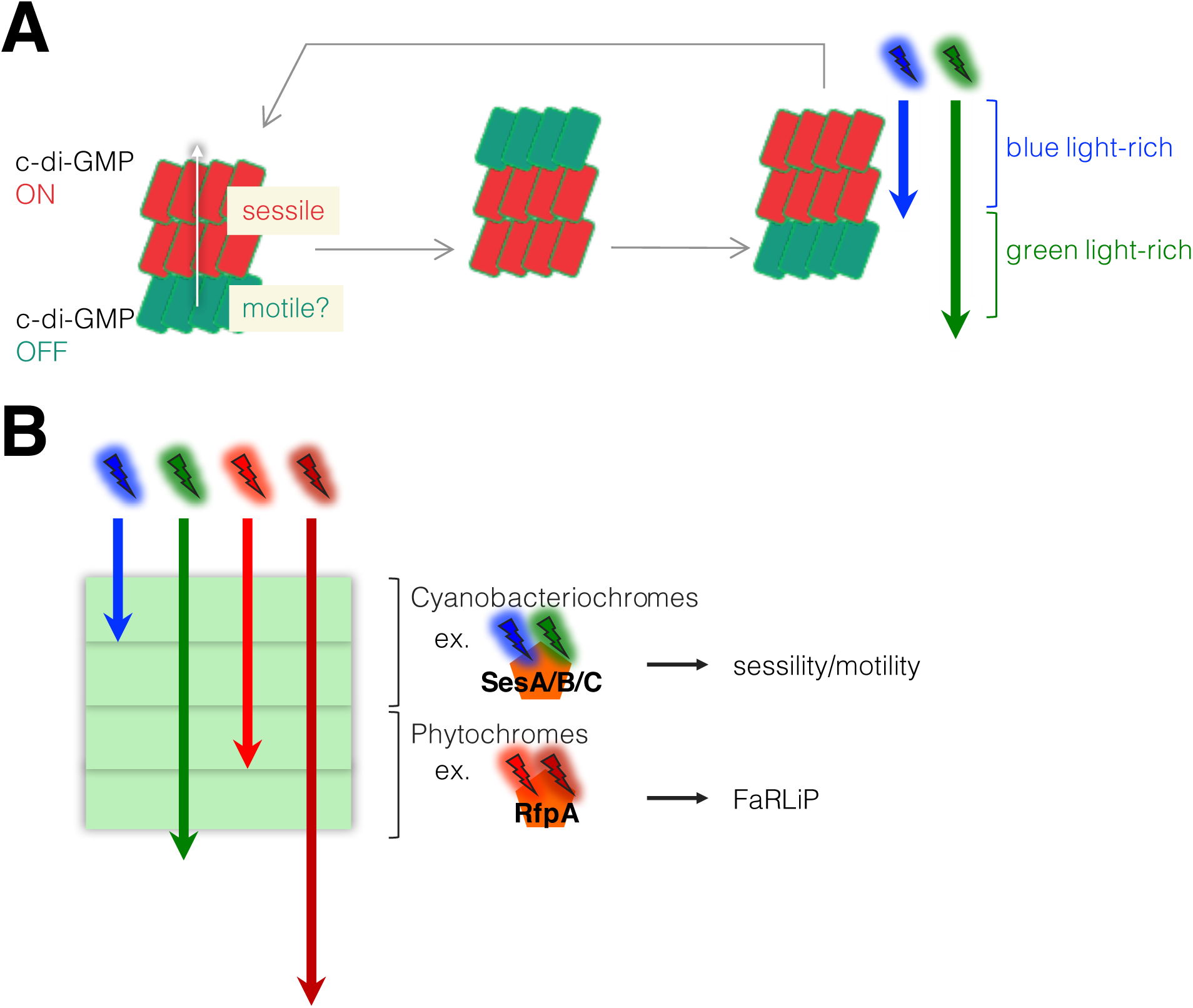
SesA/B/C Invoke Heterogeneity of C-di-GMP Signaling in a Cyanobacterial Community and Act in Different Layers than Red/Far-Red Light-Responsive Phytochromes. (A) A hypothesis of dynamic cell movements inside a microbial mat. Under natural light conditions, the internal cells in a floc will sense a green light-rich environment and turn off c-di-GMP signaling, favoring the motile-planktonic lifestyle. If the internal c-di-GMP-OFF cells retain positive phototactic motility, they may move and reach the upper region, resulting in the reorganization of the heterogenous c-di-GMP levels in the floc. This would invoke sequential cellular movements in the cyanobacterial community. (B) A hypothesis of niche differentiation of photoreceptors. Blue light rapidly attenuates in the top layer of photosynthetic microbial mats [15, 26]. Blue/green cyanobacteriochromes may be a shade detector in an upper layer of microbial mats, where sufficient red light is still available. On the other hand, phytochromes may be effective in a darker deep area, where red light diminishes, and therefore cyanobacteria cannot out-compete other bacterial species.

### Two Strategies of Light Acclimation in Mats

The phototrophic microbial mats in the Nakabusa hot spring in Japan contain unicellular *Thermosynechococcus* and chlorophyll *f*-producing filamentous cyanobacteria [15, 23]. The production of chlorophyll *f* as well as specific subunits of PSI, PSII, and phycobiliproteins for absorbing long-wavelength light is induced by far-red light via the knot-less phytochrome RfpA [1, 24, 25]. This far-red light-induced acclimation enables active photosynthesis under the red light-depleted niche when cells are embedded in other chlorophyll *a*-containing cyanobacteria. However, even these far-red light-responsive organisms, such as *Leptolyngbya* sp. JSC-1, possess proteins harboring both a blue/teal-green-type CBCR domain and domain(s) of c-di-GMP production or degradation, like SesA/B/C. This fact suggests that blue/teal-green regulation of cell aggregation (sessility) may be more widely distributed than the far-red light-induction of chlorophyll *f* and long-wavelength photosynthesis. Notably, blue light penetrates mats less than red light due the higher absorption and scattering of blue light [15, 26]. The green light may be a useful reference beam to the blue monitoring beam. The uppermost cyanobacterial layer of the microbial mats in the Nakabusa hot spring does not contain phycoerythrin or phycoerythrocyanin, which absorb green light for photosynthesis. This fact may support the idea that green light can be a reference for the recognition of differential light quality. In other words, blue/teal-green CBCRs may be superior to red/far-red phytochromes for the shade detection in the upper region of the cyanobacterial layer of microbial mats (Figure 4B) because blue and teal-green light are less penetrating than red and far-red light, respectively, in thermophilic cyanobacterial mats [15, 26]. Red/far-red phytochromes, which are well known as a shade detector for shade avoidance of land plants, are distributed in various organisms, such as plants, algae, fungi, and bacteria [3, 27], suggesting that phytochromes may be the ancestor of the related but cyanobacteria-specific photoreceptors CBCRs. The blue/green variant is one of the most prevailing features among CBCRs but is missing in any other type of photoreceptor [4, 5]. Thus, the new demand for blue/green sensors in a unique niche might be a driving force for CBCR evolution in cyanobacteria. Many blue/green CBCR variants are involved in the regulation of community-based phototactic motility [21, 28–31] and floc formation [32], suggesting that blue/green CBCRs will work as cell shade sensing systems and orchestrate motile cell communities under natural light conditions.

## Acknowledgments

This work was supported by a grant-in-aid for Young Scientists (B) (JSPS KAKENHI Grant No. 17K15244) from Japan Society for the Promotion of Science (GE) and by Core Research for Evolutional Science and Technology, Japan Science and Technology Agency (MI). GE was supported by EMBO Long-Term Fellowship (ALTF 274-2017). We thank Dr. Chihiro Azai and Prof. Dr. Satoshi Hanada for insightful discussions and suggestions. We are also grateful to Prof. Dr. Annegret Wilde and Ms. Annik Jakob for their critical reading of the manuscript.

## Author Contributions

Conceptualization, G.E. and M.I.; Methodology, G.E.; Investigation, G.E.; Writing – Original Draft, G.E.; Writing – Review & Editing, G.E. and M.I.; Funding Acquisition, G.E. and M.I.; Resources, G.E. and M.I.

## Methods

### Cell cultivation and cell aggregation assay

The thermophilic cyanobacterium *T. vulcanus* strain RKN (equivalent to National Institute for Environmental Studies 2134) that shows positive phototaxis was cultured in BG11 medium at 45°C [33]. The culture density was monitored at 730 nm. The cultures of wild-type *T. vulcanus* grew at 45°C (OD_730_ 0.5–2) and were diluted to an OD_730_ of 0.2 unless otherwise stated. These samples were then incubated at 31°C for 48 h because cell aggregation is enhanced at relatively low temperatures compared to their optimum temperatures [10, 19]. The cells were cultured under photosynthetic red light (λ_max_ = 634 nm; 30 µmol photon•m^−2^•s^−1^; Valore Corp., Kyoto, Japan) with blue or teal-green light (λ_max_ =448 or 507 nm, respectively; 5 µmol photon•m^−2^•^−1^; Valore Corp.). The cellulose-dependent cell-aggregation assay was performed as previously described, and the results are reported as an aggregation index (%) [10]. Briefly, after the irradiation period, cell suspensions were thoroughly mixed, and aliquots were transferred to cuvettes. The samples were held at room temperature in the dark for 30 min; during this time, most aggregated cells had precipitated to the bottom of the cuvettes. Then, the OD_730_ of each sample was measured (denoted OD_NA_; i.e., OD_730_ of NonAggregated cells remaining in the culture medium). Next, cellulase (12.5 U•mL^−1^; Worthington Biochemical, NJ, USA) was added to each cuvette sample, which was then incubated for 30 min at 37°C to disperse the aggregated cells completely. The OD_730_ of each sample (denoted OD_total_) was then measured. The aggregation index (%) is defined as ((OD_total_– OD_NA_)/OD_total_) × 100.

